# Ethane-dependent synthesis of polyhydroxyalkanoates by the obligate methanotroph *Methylocystis parvus* OBBP

**DOI:** 10.1101/307108

**Authors:** Myung Jaewook, James C. A. Flanagan, Wakuna M. Galega, Robert M. Waymouth, Craig S. Criddle

## Abstract

Under conditions of nutrient-limited growth, Type II obligate methanotrophs oxidize C_1_ compounds, such as methane or methanol and accumulate intracellular granules of poly(3-hydroxybutyrate) (P3HB). Here, we report that, under same nutrient-limited conditions, the Type II obligate methanotroph *Methylocystis parvus* OBBP can use ethane as its sole carbon and energy source for synthesis P3HB granules, accumulating up to 35 ± 4 wt% P3HB. ^13^C-NMR spectra of the P3HB confirmed incorporation of ^13^C from [^13^C_2_]ethane. Moreover, when valerate was added as a co-substrate with ethane, oxidation of the ethane supported synthesis of the copolymer poly(3-hydroxybutyrate-co-3-hydroxyvalerate) (PHBV).

**IMPORTANCE:** The presence of ethane in natural gas is often considered undesirable for methanotroph-based biotechnology due to the C_1_ specialization of obligate methanotrophs and concerns about inhibitory byproducts arising from methane monooxygenase-mediated cometabolism of ethane. This work establishes that co-oxidation of ethane and further metabolism in the absence of methane can support synthesis of the valuable polyhydroxyalkanoate bioplastics P3HB and PHBV.

## INTRODUCTION

Aerobic methanotrophs are a unique group of gram-negative bacteria capable of utilizing methane as sole carbon and energy source (1). The control of methanotrophs is critical as they play a key role in mitigating the greenhouse gas impacts of methane (2) and can produce nitrous oxide, an even more powerful global warming agent (3). Biotechnological applications of methanotrophic bacteria include production of biodiesel (4), propylene oxide (5), single cell protein (6), extracellular polysaccharides (7), human health supplements (8), and bioplastics, including poly(3-hydroxybutyrate) (P3HB) (9–11), poly(3-hydroxybutyrate-*co*-3-hydroxyvalerate) (PHBV) (11–14), poly(3-hydroxybutyrate-*co*-4-hydroxybutyrate), poly(3-hydroxybutyrate-*co*-5-hydroxyvalerate-*co*-3-hydroxyvalerate), and poly(3-hydroxybutyrate-*co*-6-hydroxyhexanoate-*co*-4-hydroxybutyrate) (15).

Ethane (C_2_H_6_) is of interest for methanotroph-based biotechnology because it is the second most common component of natural gas (up to 5-10%) after methane (16). Because the obligate methanotrophs are one-carbon specialists, ethane is typically depicted as an undesirable contaminant (17). While cometabolic oxidation of ethane to ethanol by methane monoxygenase is well-known, soluble products resulting from this oxidation are often viewed as inhibitory and an obstacle to beneficial use of natural gas. In this study, we report P3HB production by a pure culture of *Methylocystis parvus* OBBP when ethane is added as the sole source of carbon during nitrogen-limited, unbalanced growth. We further demonstrate that ethane oxidation can support synthesis of PHBV when valerate is provided as a co-substrate during unbalanced growth.

## MATERIALS AND METHODS

### Culture conditions

Unless otherwise specified, all *Methylocystis parvus* OBBP cultures were grown in medium JM2, which is a modified version of ammonium mineral salts (AMS) medium (18). Medium JM2 contained the following chemicals per L of solution: 2.4 mM MgSO_4_ · 7H_2_O, 26 mM CaCl_2_, 36 mM NaHCO_3_, 4.8 mM KH_2_PO_4_, 6.8 mM K_2_HPO_4_, 10.5 μM Na_2_MoO_4_ · 2H_2_O, 7 μM CuSO_4_ · 5H_2_O, 200 μM Fe-EDTA, 530 μM Ca-EDTA, 5 mL trace metal solution, and 20 mL vitamin solution. The trace stock solution contained the following chemicals per L of solution: 500 mg FeSO_4_ · 7H_2_O, 400 mg ZnSO_4_ · 7H_2_O, 20 mg MnCl_2_ · 7H_2_O, 50 mg CoCl_2_ · 6H_2_O, 10 mg NiCl_2_ · 6H_2_O, 15 mg H_3_BO_3_ and 250 mg EDTA. The vitamin stock solution contained the following chemicals per L of solution: 2.0 mg biotin, 2.0 mg folic acid, 5.0 mg thiamine · HCl, 5.0 mg calcium pantothenate, 0.1 mg vitamin B12, 5.0 mg riboflavin and 5.0 mg nicotinamide.

All cultures were incubated in 160 mL serum bottles (Wheaton, Millville, NJ, USA) capped with butyl-rubber stoppers and crimp-sealed under a methane/oxygen headspace (molar ratio 1:1.5; > 99% purity; Praxair Technology, Inc., Danbury, CT, USA). Liquid volume was 50 mL, and the headspace volume was 110 mL. Cultures were incubated horizontally on orbital shaker tables at 150 rpm. The incubation temperature was 30 °C.

### Methane-fed balanced growth phase

Fifty-milliliter cultures were grown to final optical densities (OD_600_) of 0.8-1.2 then centrifuged (3000 × g) for 15 min. The pellets were resuspended in 30 mL of JM medium to create the inoculum for triplicate 160 mL serum bottle cultures. Each culture received 10 mL inoculum plus 40 mL of fresh medium (39.5 mL of medium JM plus 0.5 mL of 1.35 M ammonium chloride stock) and was flushed for 5 min with a methane/oxygen mixture (molar ratio of 1:1.5). After growth at 30 °C for 24 h, the headspace in each culture was again flushed for 5 min with the methane/oxygen mixture then incubated at 30 °C for a second 24 h period of balanced growth. All experiments were carried out in triplicate.

### Ethane-fed unbalanced growth phase

After 48 h, all cultures were harvested and subjected to nitrogen-limiting conditions. Triplicate samples were centrifuged (3000 × g) for 15 min and suspended in fresh medium without nitrogen. The headspace of each bottle was flushed with the ethane/oxygen gas mixture (molar ratio of 1:1.5, > 99% purity; Praxair Technology, Inc., Danbury, CT, USA) at t = 0 h and t = 24 h. To confirm ethane incorporation into the P3HB granules, [^13^C_2_]ethane (99 atom% ^13^C_2_ ethane, Sigma-Aldrich, St. Louis, MO, USA) was added in some cycles.

### Ethane-fed unbalanced growth phase plus valerate

After detection of P3HB from ethane-growth cell cultures, tests were performed to determine whether ethane could support production of PHBV when valerate was added during the PHA accumulation phase. Grown cells were centrifuged (3500 rpm) for 15 min to create a pellet. The cell pellet was re-suspended in 50 mL of JM2 media without added nitrogen, and vortexed vigorously to obtain a uniform suspension. The suspension was transferred into 160-mL serum bottle, capped and crimp sealed. Valerate (10 mM) was added as sodium valerate to a subset of the serum bottles to determine whether ethane could support production of PHBV. The headspace was flush with C_2_H_6_:O_2_ mixture (molar ratio 1:4; > 99% purity; Praxair Technology, Inc., Danbury, CT, USA) and incubated for 48 hr. At the 24-hr midway point, the headspace of each vial was flushed with the same ethane mixture. At the end of the 48-hr PHA accumulation period, the cells were harvested, centrifuged (3000 × g) for 15 min to create a pellet, and then freeze dried for further PHA analysis.

### Ethane oxidation products analysis

In some ethane-fed unbalanced growth phases, 1 mL of cell suspensions were sampled every 15 min during the first initial hour to analyze the concentrations of oxidation products (alcohols, aldehydes, and carboxylic acids) of ethane. Products were determined using gas chromatography (detailed methods are discussed in Analytical methods).

### Culture purity check

To test culture purity, biomass was removed after the 48 h period of balanced growth. Genomic DNA was extracted using the FastDNA SPIN Kit for Soil (MP Biomedicals, Santa Ana, CA, USA), as per the manufacture’s protocol. Bacterial 16S rRNA was amplified using the bacterial primers BAC-8F (5’-AGAGTTTGATCCTGGCTCAG-3’) and BAC-1492R (5’-CGGCTACCTTGTTACGACTT-3’) (19). A polymerase chain reaction (PCR) was performed using AccuPrime Taq DNA Polymerase System (Invitrogen, Carlsbad, CA, USA) with the following thermocycling steps: (i) 94 °C for 5 min; (ii) 30 cycles consisting of 94 °C for 30 s, 55 °C for 30 s, 68 °C for 80 s; and (iii) an extension at 68 °C for 10 min. Amplicon presence and quality of PCR reaction were verified via 1.5% agarose gel electrophoresis.

PCR products were purified using QIAquick PCR Purification Kit (Qiagen, Chatsworth, CA, USA), then cloned using pGEM-T Easy Vector System with JM109 competent Escherichia coli cells (Promega, Madison, WI, USA) per the manufacture’s protocol. Randomly selected clones were sequenced by Elim Biopharmaceuticals Inc. (Hayward, CA, USA), generating 120 near-full length 16S rRNA gene sequences. Retrieved DNA sequences were compared with reference sequences using Basic Local Alignment Search Tool (BLAST).

### Confirmation of obligate methanotrophy

To test use of ethane as a carbon source for growth, cultures were incubated under ethane/oxygen gas mixture (molar ratio of 1:1.5, > 99% purity; Praxair Technology, Inc., Danbury, CT, USA) in 50 mL of fresh medium (49.5 mL of medium JM plus 0.5 mL of 1.35 M ammonium chloride stock). Control cultures without any added carbon source were also prepared. To rule out the effect of O_2_ tension in methanotrophic growth, various concentrations of O_2_ (1, 5, 10, and 25% O_2_ in the headspace) were tested. OD_600_ values were measured for 10 days.

### Analytical methods

To analyze concentrations of methane, ethane, and oxygen, 0.5 mL of gas phase from each enrichment culture was injected onto GOW-MAC gas chromatograph with an Altech CTR 1 column and a thermal conductivity detector. The following method parameters were used: injector, 120 °C; column, 60 °C; detector, 120 °C; and current, 150 mV. Peak areas of methane, ethane, and oxygen were compared to standards and quantified using the software ChromPerfect (Justice Laboratory Software, Denville, NJ, USA).

Products of ethane oxidation (alcohols and aldehydes) were analyzed using a GC (Agilent 6890N; Agilent Technologies, Palo Alto, CA, USA) equipped with an HP-INNOWax column (Agilent Technologies, Palo Alto, CA, USA) and a flame ionization detector.

To analyze total suspended solids (TSS), 0.5-5.0 mL of cell suspension was filtered through prewashed, dried, and pre-weighted 0.2 μm membrane filters (Pall, Port Washington, NY, USA).

The filtered cells and membrane filters were dried at 105 °C for 24 h, then weighed on an AD-6 autobalance (PerkinElmer, Norwalk, CT, USA).

### PHA weight percentages

To determine PHA weight percent, between 5 and 10 mg of freeze-dried biomass were weighed then transferred to 12 mL glass vials. Each vial was amended with 2 mL of methanol containing sulfuric acid (3%, vol/vol) and benzoic acid (0.25 mg/mL methanol), supplemented with 2 mL of chloroform, and sealed with a Teflon-lined plastic cap. All vials were shaken then heated at 95—100 °C for 3.5 h. After cooling to room temperature, 1 mL of deionized water was added to create an aqueous phase separated from the chloroform organic phase. The reaction cocktail was mixed on a vortex mixer for 30 s then allowed to partition until phase separation was complete. The organic phase was then sampled by syringe and analyzed using a GC (Agilent 6890N; Agilent Technologies, Palo Alto, CA, USA) equipped with an HP-5 column (containing (5% phenyl)-methylpolysiloxane; Agilent Technologies, Palo Alto, CA, USA) and a flame ionization detector. DL-3-Hydroxybutyric acid sodium salt (Sigma-Aldrich, St. Louis, MO, USA) and PHBV with 3HV fractions of 5 mol%, 8 mol%, and 12 mol% (Sigma-Aldrich, St Louis, MO, USA) was used to prepare external calibration curves. The PHA content (wt%, w_PHA_/w_CDW_) of the samples was calculated by normalizing to initial dry mass.

### PHA purification

PHA was quantified using the gas chromatography protocol of Braunegg et al. (20). PHA granules were extracted from the cells by suspending 500 mg of freeze-dried cell material in 50 mL Milli-Q water, adding 400 mg of sodium dodecyl sulfate (>99.0% purity; Sigma-Aldrich, St. Louis, MO, USA) and 360 mg of EDTA, followed by heating to 60 °C for 60 min to induce cell lysis. The solution was centrifuged (3000 × g) for 15 min, and the pellet washed three times with deionized water. To purify the PHA, pellets were washed with a 50 mL sodium hypochlorite (bleach) solution (Clorox 6.15%), incubated at 30 °C with continuous stirring for 60 min, then centrifuged (3000 × g) for 15 min. Sample pellets were washed and recentrifuged three times with deionized water.

### Molecular weight analysis

Molecular weights of PHAs were evaluated using gel permeation chromatography (GPC). Sample pellets dissolved in chloroform at a concentration of 5 mg/mL for 90 min at 60 °C were filtered through a 0.2 μm PTFE filter, then analyzed with a Shimadzu UFLC system (Shimadzu Scientific Instruments, Columbia, MD, USA) equipped with a Shimadzu RID-10A refraction index detector. The GPC was equipped with a Jordi Gel DVB guard column (500 Å, Jordi Labs, Mansfield, MA, USA) and Jordi Gel DVB analytical columns (105 Å, Jordi Labs, Mansfield, MA, USA). The temperature of the columns was maintained at 40 °C, and the flow rate of the mobile phase (chloroform) was 1 mL min^-1^. Molecular weights were calibrated with polystyrene standards from Varian (Calibration Kit S-M2-10, Agilent Technologies, Palo Alto, CA, USA).

### Nuclear magnetic resonance (NMR)

NMR spectroscopy was used to detect the abundance of ^13^C in P3HB samples made from bacteria fed either naturally-abundant (99% ^12^C, 1% ^13^C) (control sample) or isotopically-labeled (99% ^13^C, 1% ^12^C) ethane. A stock solution of internal standard was made by dissolving 1.0 mg hexamethyldisiloxane (HMDSO) in 1.44 mL CDCl_3_. For both samples, 0.5 mL of this internal standard stock solution (2.130 x 10^−3^ mmol) was combined with approximately 1 mg of P3HB (an accurate amount of P3HB in the sample was determined by ^1^H-NMR: *vide infra*). ^1·^H-NMR spectroscopy (500 MHz, 32 scans, delay time (d1) = 20 s) was firstly used to calculate an accurate amount of dissolved P3HB in the samples by comparing peak integrations of the CH_3_ protons from HMDSO and CH_3_ protons from P3HB. ^13^C-NMR spectroscopy (125 MHz, 2044 scans (for P3HB from naturally abundant ethane) and 2392 scans (for P3HB from [^13^C_2_]-ethane), delay time (d1) = 16 s, room temperature, 90° pulse) was then performed on the two samples. In both samples, a comparison of the peak integrations of the CH_3_ carbons from HMDSO and CH_3_ carbon from P3HB was used to determine whether the carbon was present in a natural abundance or in greater than natural abundance (which would indicate incorporation of ^13^C-carbons from the [^13^C_2_]-ethane).

## RESULTS

### Culture purity check and tests of growth with ethane

Culture purity checks gave no indication of culture contamination by heterotrophic bacteria (21, 22). All cloned 16S rRNA gene fragments matched reference sequences for *M. parvus* OBBP. This strain was tested for growth on ethane. No growth was detected on ethane over a range of headspace O_2_ levels (1, 5,10 and 25%), confirming obligate methanotrophy.

### P3HB production using ethane

Table 1 summarizes P3HB production results for *M. parvus* OBBP when methane or ethane was added as a carbon source during a nitrogen-limited, unbalanced growth phase. In either case, P3HB was the final product. When methane was added, the wt% P3HB was 48 ± 5 wt%, and the TSS was 1820 ± 200 mg/L. When ethane was added, the wt% P3HB decreased to 35 ± 4 wt%, and the TSS decreased to 1440 ± 160 mg/L.

**Table 1.** Final P3HB wt% and TSS values after a 48 h nitrogen-limited, unbalanced growth phase when methane or ethane was added as a carbon source.

| Carbon source | P3HB wt% | TSS (mg/L) |
| --- | --- | --- |
| Methane | $48 \pm 5$ | $1820 \pm 200$ |
| Ethane | $35 \pm 4$ | $1440 \pm 160$ |

Patterns of gas consumption and generation were evaluated for the serum bottle cultures fed methane (Figure 1a) or ethane (Figure 1b) on a nitrogen-limited, unbalanced growth phase. The errors bars represent standard deviations for triplicate batch cultures.

**Figure 1.**
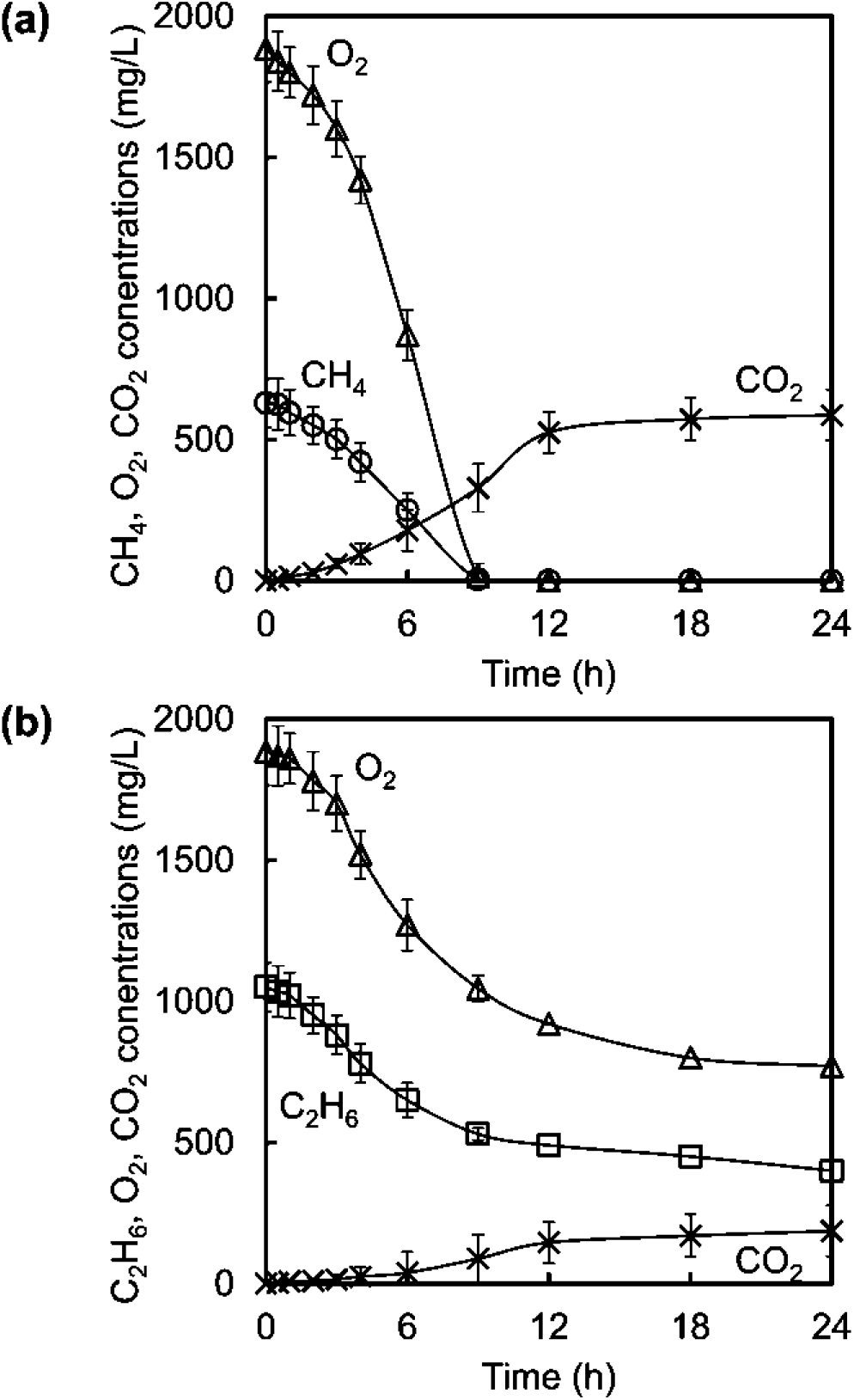
Concentrations of methane 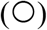, ethane 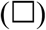, oxygen 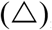, and carbon dioxide 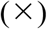 measured during the nitrogen-limited, unbalanced growth phase when (a) methane or (b) ethane was added as a carbon source. Gases in both the gas phase and the liquid phase were taken account to compute for the concentrations. The errors bars represent standard deviations for triplicate enrichment cultures.

When fed methane, almost all methane and oxygen were consumed within the first 9 h. The final concentration of CO_2_ was 587 ± 90 mg CO_2_/L. The maximum specific rate of methane utilization (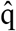_CH4_) was 0.058 ± 0.009 g CH_4_ g TSS^-1^ h^-1^. The maximum specific rate of oxygen utilization (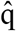_O2_) was 0.164 ± 0.021 g O_2_ g TSS^-1^ h^-1^. The maximum specific rate of CO_2_ production (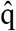_CO2_) was 0.038 ± 0.007 g CO_2_ g TSS^-1^ h^-1^. No inhibition was observed throughout the 24 h period.

When fed ethane, both ethane and oxygen were present throughout the 24 h period. The final concentration of CO_2_ was 185 ± 33 mg CO_2_/L. The maximum specific rate of ethane utilization(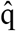_C2H6_) was 0.048 ± 0.008 g C_2_H_6_ g TSS^-1^ h^-1^. The maximum specific rate of oxygen utilization (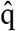_O2_) was 0.088 ± 0.015 g O_2_ g TSS^-1^ h^-1^. The maximum specific rate of CO_2_ production (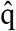_CO2_) was 0.010 ± 0.002 g CO_2_ g TSS^-1^ h^-1^. The consumption rate of ethane and oxygen slowed down after t = 6 h, suggesting the presence of inhibitory substances in the cell cultures.

### Isotopic enrichment

^13^C-NMR spectroscopy was used to detect the abundance of ^13^C in P3HB samples made from bacteria fed either naturally-abundant (99% ^12^C, 1% ^13^C) (control sample) or isotopically-labeled (99% ^13^C, 1% ^12^C) ethane. A HMDSO internal standard was used in both samples (see full Supplemental Material for full calculation). In the control sample, where the P3HB was synthesized from bacteria supplied with naturally abundant ethane, the carbon peak integrals indicated that the ^13^C-content in the polymer was approximately 1% (i.e. consistent with the 1% natural abundance of ^13^C) (Figure 2b). In contrast, in the P3HB sample from bacteria supplied with [^13^C_2_]-ethane, the carbon peak integrals indicated that the ^13^C-content in the polymer was approximately 10% (Figure 3b). This demonstrates unequivocally that the bacteria uptake ^13^C-labeled ethane, and that it is utilized in the synthesis of P3HB.

**Figure 2.**
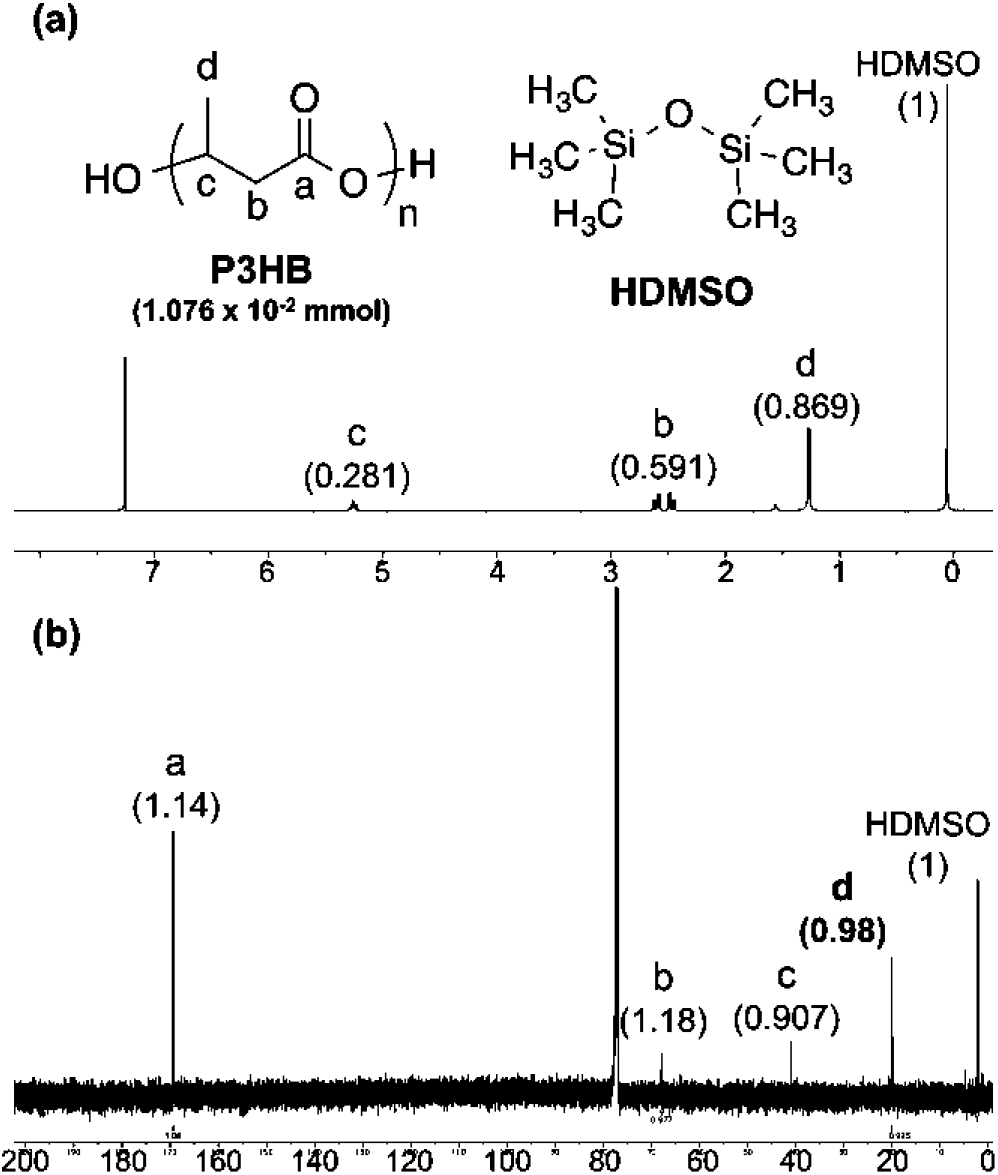
(a) ^1^H-and (b) ^13^C-NMR spectra of P3HB polymer produced using naturally abundant ethane. Numbering of the atoms is illustrated on a chemical structure. Numbers inside the parentheses are the corresponding peaks’ integration numbers.

**Figure 3.**
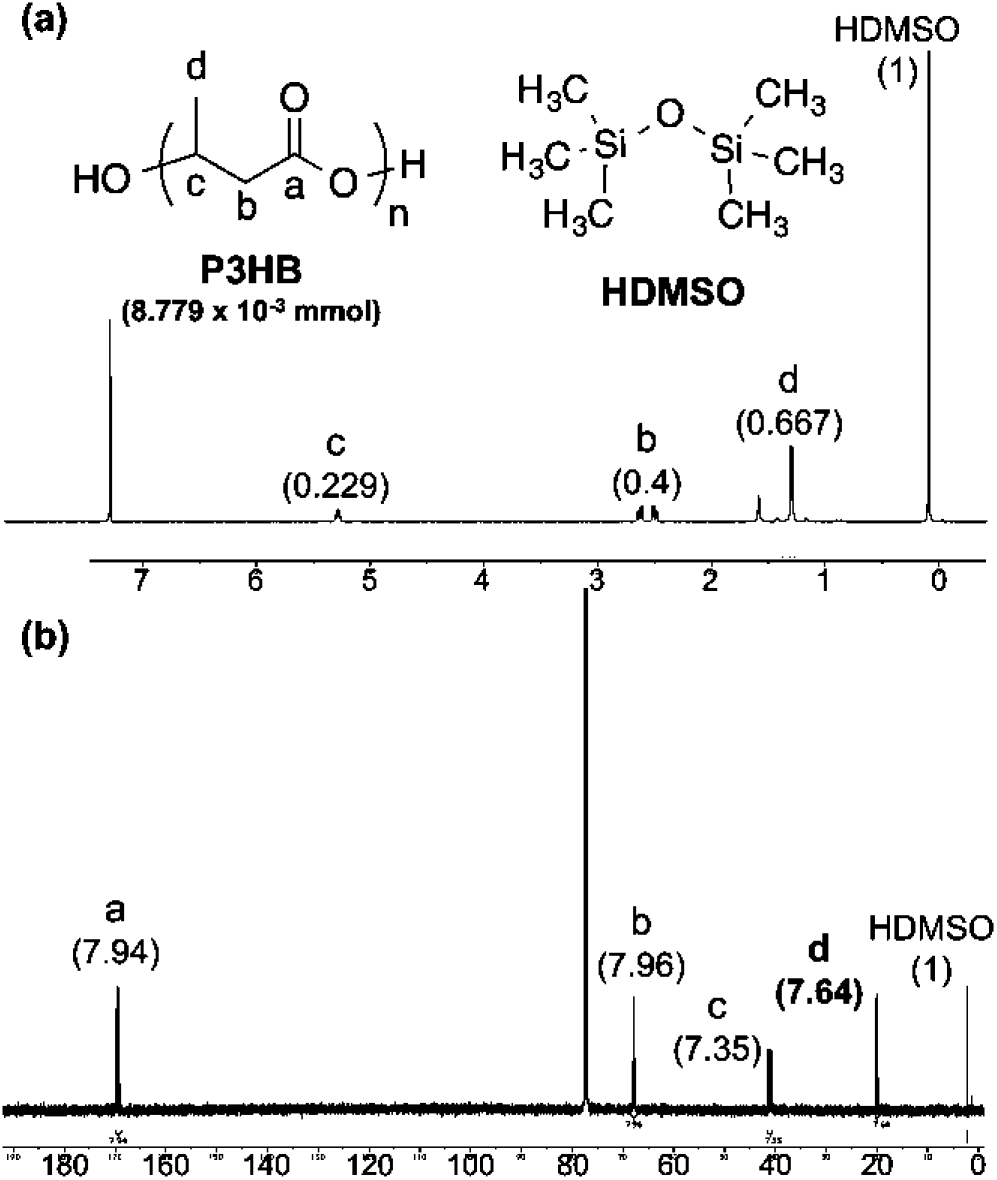
(a) ^1^H-and (b) ^13^C-NMR spectra of P3HB polymer produced using [^13^C_2_]ethane. Numbering of the atoms is illustrated on a chemical structure. Numbers inside the parentheses are the corresponding peaks’ integration numbers.

### Molecular weight characterization

Table 2 illustrates the number average molecular weight (M_n_) and molecular weight distributions (M_w_/M_n_) of P3HB produced by *M.parvus* OBBP when fed methane or ethane. Values for M_n_ and M_w_/M_n_ are not statistically different when fed methane or ethane (p-value of M_n_ = 0.75, p-value of M_w_/M_n_ = 0.43).

**Table 2.**
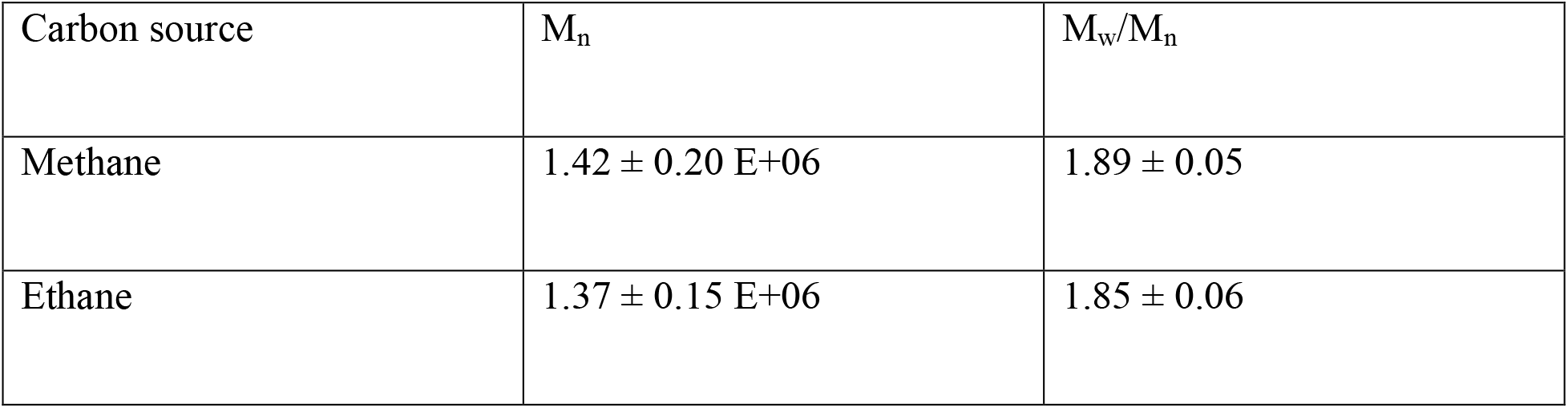
Molecular weights and distributions for extracted P3HB.

| Carbon source | $M_n$ | $M_w/M_n$ |
| --- | --- | --- |
| Methane | $1.42 \pm 0.20 \text{ E}+06$ | $1.89 \pm 0.05$ |
| Ethane | $1.37 \pm 0.15 \text{ E}+06$ | $1.85 \pm 0.06$ |

### Oxidation products of ethane

Cell cultures of *M. parvus* OBBP oxidized ethane gas. Products from the oxidation of ethane included acetaldehyde and acetate (Table 3). Ethanol was not detected, suggesting that *M. parvus* OBBP has an efficient alcohol dehydrogenase system.

**Table 3.** Oxidation products of ethane by cell cultures of *M. parvus* OBBP during the first hour in a nitrogen-limited, unbalanced growth phase

| Products detected | Rate of formation (mg product mg TSS<sup>-1</sup> h<sup>-1</sup>) |
| --- | --- |
| Ethanol | ND |
| Acetaldehyde | 0.066 ± 0.011 |
| Acetate | 0.71 ± 0.13 |

### PHBV production from ethane and valerate

When ethane was the source of carbon and energy during the PHA accumulation phase, P3HB was the sole PHA produced; when supplemented with valerate (10 mM), 12.9 ± 2.6% of PHBV was generated (25 ± 0.01% 3HV mole fraction).

## DISCUSSION

Obligate methanotrophs are restricted to grow on C_1_ substrates including methane, methanol, and in some cases formate, formaldehyde, and methylamines (23). While *M. parvus* OBBP is unable to grow on ethane under conditions of balanced, nutrient-sufficient growth, this work establishes that *M. parvus* OBBP is able to take up ethane and oxidized it under nitrogen-limited, unbalanced growth conditions. Methane monooxygenases (MMOs) are known to be non-specific (17, 24) and capable of oxidizing aliphatic compounds, aromatic compounds, and alkanes, including ethane (25–31), and, our data indicate that MMO-generated ethanol degrades rapidly to acetaldehyde then acetic acid. A fraction of the acetic acid is assimilated into P3HB. This is the first evidence that pure culture of well-known Type II obligate methanotrophs assimilate ethane and produce P3HB polymer with high molecular weight comparable to the P3HB polymer made with methane (Table 2). Incorporation of ^13^C-labeled ethane was confirmed using ^13^C-NMR.

Figure 4 illustrates the proposed pathway for oxidation and assimilation of ethane. All the presented enzymes in *M. parvus* OBBP have been identified and their primary structures are deposited in publically available databases.

**Figure 4.**
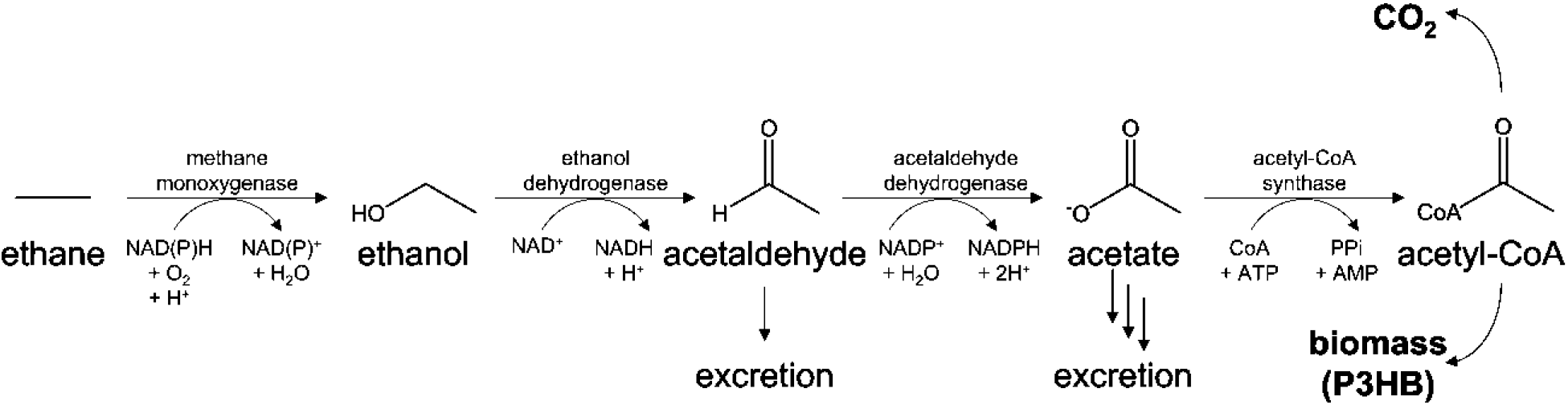
Pathway for assimilation of ethane by *M. parvus* OBBP.

MMO oxidation of ethane to ethanol requires reducing equivalents, which can be obtained from each subsequent oxidation step. Oxidation of ethane yielded mixtures of intermediates in the medium including acetaldehyde and acetate, but no ethanol (Table 3). The absence of ethanol and low levels of acetaldehyde suggest that *M. parvus* OBBP possesses a highly efficient dehydrogenase system.

Our data indicate that acetic acid is produced inside intracellularly by oxidation of ethane to ethanol and the acid generated is secreted into solution. The rates of intracellular oxidation of ethanol to acetate exceed the rates of acetic acid oxidation and assimilation into P3HB, resulting in the secretion of acetic acid and its accumulation in the media. Levels of acetate in the media were much higher rate than those of acetaldehyde (Table 3), in close agreement with that reported previously (31). This result and our analysis of P3HB production (Table 1) indicate that *M. parvus* OBBP possesses the enzyme systems needed to funnel acetic acid into the P3HB biosynthetic pathway, with the balance either excreted or oxidized for energy. In the presence of ethane, MMO activity decreased over time, perhaps due to dissipation of proton gradient needed for ATP production as acetic acid accumulated (Figure 1b) (32, 33).

In summary, the results from this study establish that pure cultures of obligate methanotrophs can process the two most dominant gases in natural gas into P3HB and PHBV without methane. This may be industrially significant given that ethane is typically the second-largest component of natural gas. Our findings also have implications for the way in which we view the methanotrophic P3HB biosynthesis. Until now, researchers have largely assumed that obligate methanotrophs are limited to utilization of C_1_ compounds. This work demonstrates that this basic assumption is incorrect for nutrient-limited, unbalanced growth conditions. Under such conditions, *M. parvus* OBBP can use at least two gas substrates as well as multi-carbon cosubstrates for production of PHAs. Of course, additional studies are needed to generalize this finding for other Type II methanotrophs. In preliminary tests of *Methylosinus trichosporium* OB3b, ethane addition did not support P3HB production (data not shown).

## ACKNOWLEDGEMENTS

This work was supported in part by unrestricted gifts from Chevron and by a Samsung Scholarship. This work was also supported by the Center for the Utilization of Biological Engineering in Space (CUBES), a Space Technology Research Institute grant from NASA’s Space Technology Research Grants Program under grant or cooperative agreement award number NNXNNX17AJ31G S02. We thank the Stanford Nano Shared Facilities for staff assistance, training, and access to instruments required for this research. JF is grateful to the Stanford Center for Molecular Analysis and Design (CMAD) for a graduate fellowship.

## REFERENCES

1. Karthikeyan OP, Karthigeyan K, Cirés S, Heimann K. 2014. Review of sustainable methane mitigation and biopolymer production. Crit Rev Environ Sci Technol 45:1579–1610.

2. Singh BK, Bardgett RD, Smith P, Reay DS. 2010. Microorganisms and climate change: terrestrial feedbacks and mitigation options. Nat Rev Microbiol 8:779–790.

3. Myung J, Wang Z, Yuan T, Zhang P, Van Nostrand JD, Zhou J, Criddle CS. 2015. Production of nitrous oxide from nitrite in stable Type II methanotrophic enrichments. Environ Sci Technol 49:10969–10975.

4. Fei Q, Guarnieri MT, Tao L, Laurens LML, Dowe N, Pienkos PT. 2014. Bioconversion of natural gas to liquid fuel: Opportunities and challenges. Biotechnol Adv 32:596–614.

5. Hou C. 1984. Propylene oxide production from propylene by immobilized whole cells of Methylosinus sp. CRL 31 in a gas-solid bioreactor. Appl Microbiol Biotechnol 19:1–4.

6. Yazdian F, Hajizadeh S, Shojaosadati SA, Khalilzadeh R, Jahanshahi M, Nosrati M. 2005. Production of single cell protein from natural gas: parameter optimization and RNA evaluation. Iran J Biotechnol 3:235–242.

7. Chiemchaisri W, Wu JS, Visvanathan C. 2001. Methanotrophic production of extracellular polysaccharide in landfill cover soils. Water Sci Technol 43:151–158.

8. Müller H, Hellgren LI, Olsen E, Skrede A. 2004. Lipids rich in phosphatidylethanolamine from natural gas-utilizing bacteria reduce plasma cholesterol and classes of phospholipids: A comparison with soybean oil. Lipids 39:833–841.

9. Wendlandt KD, Jechorek M, Helm J, Stottmeister U. 2001. Producing poly-3-hydroxybutyrate with a high molecular mass from methane. J Biotechnol 86:127–133.

10. Myung J, Kim M, Pan M, Criddle CS, Tang SKY. 2016. Low energy emulsion-based fermentation enabling accelerated methane mass transfer and growth of poly(3-hydroxybutyrate)-accumulating methanotrophs. Bioresour Technol 207:302–307.

11. Flanagan JCAJCA, Myung J, Criddle CSCS, Waymouth RMRM. 2016. Poly(hydroxyalkanoate)s from waste biomass: A combined chemical-biological approach. ChemistrySelect 1:2327–2331.

12. Myung J, Flanagan JCA, Waymouth RM, Criddle CS. 2016. Methane or methanol-oxidation dependent synthesis of poly(3-hydroxybutyrate-*co*-3-hydroxyvalerate) by obligate Type II methanotrophs. Process Biochem 51:561–567.

13. Cal AJ, Sikkema WD, Ponce MI, Franqui-Villanueva D, Riiff TJ, Orts WJ, Pieja AJ, Lee CC. 2016. Methanotrophic production of polyhydroxybutyrate-*co*-hydroxyvalerate with high hydroxyvalerate content. Int J Biol Macromol 87:302–307.

14. Myung J, Galega WM, Van Nostrand JD, Yuan T, Zhou J, Criddle CS. 2015. Long-term cultivation of a stable Methylocystis-dominated methanotrophic enrichment enabling tailored production of poly(3-hydroxybutyrate-*co*-3-hydroxyvalerate). Bioresour Technol 198:811–818.

15. Myung J, Flanagan JCA, Waymouth RM, Criddle CS. 2017. Expanding the range of polyhydroxyalkanoates synthesized by methanotrophic bacteria through the utilization of omega-hydroxyalkanoate co-substrates. AMB Express 7:118.

16. Mueller JC. 1969. Preferential utilization of the methane component of natural gas by a mixed culture of bacteria. Can J Microbiol 15:1114–1116.

17. Malashenko Y, Sokolov I, Romanovskaya V. 2000. Role of monooxygenase reaction during assimilation of non-growth substrates by methanotrophs. J Mol Catal-B Enzym 10:305–312.

18. Whittenbury R, Phillips KC, Wilkinson JF. 1970. Enrichment, isolation and some properties of methane-utilizing bacteria. J Gen Microbiol 61:205–218.

19. Stackebrandt E, Goodfellow M. 1991. Nucleic acid techniques in bacterial systematics. Wiley, Chichester; New York.

20. Braunegg G, Sonnleitner B, Lafferty RM. 1978. Rapid Gas-Chromatographic Method for Determination of Poly-Beta-Hydroxybutyric Acid in Microbial Biomass. Eur J Appl Microbiol Biotechnol 6:29–37.

21. Dedysh SN, Dunfield PF. 2011. Facultative and obligate methanotrophs: How to identify and differentiate them. Methods Enzymol 495:31–44.

22. Semrau JD, DiSpirito AA, Vuilleumier S. 2011. Facultative methanotrophy: false leads, true results, and suggestions for future research. FEMS Microbiol Lett 323:1–12.

23. Dedysh SN, Knief C, Dunfield PF. 2005. Methylocella species are facultatively methanotrophic. J Bacteriol 187:4665–4670.

24. Hou CT, Patel RN, Laskin AI, Barnabe N. 1980. Microbial oxidation of gaseous hydrocarbons: Oxidation of lower N-alkenes and N-alkanes by resting cell suspensions of various methylotrophic bacteria, and the effect of methane metabolites. FEMS Microbiol Lett 9:267–270.

25. Criddle CS. 1993. The kinetics of cometabolism. Biotechnol Bioeng 41:1048–1056.

26. Alvarez-Cohen L, McCarty PL. 1991. Product toxicity and cometabolic competitive inhibition modeling of chloroform and trichloroethylene transformation by methanotrophic resting cells. Appl Environ Microbiol 57:1031–1037.

27. Fogel MM, Taddeo AR, Fogel S. 1986. Biodegradation of chlorinated ethenes by a methane-utilizing mixed culture. Appl Environ Microbiol 51:720–724.

28. Higgins IJ, Best DJ, Hammond RC. 1980. New findings in methane-utilizing bacteria highlight their importance in the biosphere and their commercial potential. Nature 286:561–564.

29. Oldenhuis R, Vink RLJM, Janssen DB, Witholt B. 1989. Degradation of chlorinated aliphatic hydrocarbons by Methylosinus trichosporium OB3b expressing soluble methane monooxygenase. Appl Environ Microbiol 55:2819–2826.

30. Patel RN, Hou CT, Laskin AI, Felix A. 1982. Microbial oxidation of hydrocarbons: Properties of a soluble methane monooxygenase from a facultative methane-utilizing organism, Methylobacterium sp. strain CRL-26. Appl Environ Microbiol 44:1130–1137.

31. Imai T, Takigawa H, Nakagawa S, Shen G-J, Kodama T, Minoda Y. 1986. Microbial oxidation of hydrocarbons and related compounds by whole-cell suspensions of the methane-oxidizing bacterium H-2. Appl Environ Microbiol 52:1403–1406.

32. Axe DD, Bailey JE. 1995. Transport of lactate and acetate through the energized cytoplasmic membrane of Escherichia coli. Biotechnol Bioeng 47:8–19.

33. Yoon S, Im J, Bandow N, Dispirito AA, Semrau JD. 2011. Constitutive expression of pMMO by Methylocystis strain SB2 when grown on multi-carbon substrates: Implications for biodegradation of chlorinated ethenes. Environ Microbiol Rep 3:182–188.

